# Non-specific effects of a CINNAMATE-4-HYDROXYLASE inhibitor on auxin homeostasis

**DOI:** 10.1101/2021.10.27.466161

**Authors:** Ilias El Houari, Petr Klíma, Alexandra Baekelandt, Paul Staswick, Veselina Uzunova, Charo I. Del Genio, Ward Steenackers, Petre I. Dobrev, Ondrej Novák, Richard Napier, Jan Petrášek, Dirk Inzé, Wout Boerjan, Bartel Vanholme

## Abstract

Chemical inhibitors are often implemented for the functional characterization of genes to overcome the limitations associated with genetic approaches. Although being a powerful tool, off-target effects of these inhibitors are easily overlooked in a complex biological setting. Here we illustrate the implications of such secondary effects by focusing on piperonylic acid (PA), an inhibitor of CINNAMATE-4-HYDROXYLASE (C4H) that is often used to investigate the involvement of lignin during plant growth and development. When supplied to plants, we found that PA is recognized as a substrate by GRETCHEN HAGEN 3.6 (GH3.6), an amido synthetase involved in the formation of the auxin catabolite indole-3-acetic acid (IAA)-Asp. By competing for the same enzyme, PA interferes with auxin conjugation, resulting in an increase in cellular auxin concentrations. These increased auxin levels likely further contribute to an increase in adventitious rooting previously observed upon PA-treatment. Despite the focus on GH3.6 in this report, PA is conjugated by an array of enzymes and their subsequent reduced activity on native substrates could potentially affect a whole set of physiological processes in the plant. We conclude that surrogate occupation of the endogenous conjugation machinery in the plant by exogenous compounds is likely a more general phenomenon that is rarely considered in pharmacological studies. Our results hereby provide an important basis for future reference in studies using chemical inhibitors.

## INTRODUCTION

Unraveling the physiological function of genes is challenging and a frequent strategy towards this goal is the use of loss-of-function mutants. Such strategies however come with certain limitations. Due to gene redundancy or compensation mechanisms, phenotypes can for instance be masked and if lethal phenotypes are obtained further analysis of the mutants is severely hampered (Bouché and Bouchez 2001; Rohde et al. 2004; El Houari et al. 2021b). An alternative approach is to use chemical inhibitors to interfere with the protein of interest and mimic loss-of-function mutants. These inhibitors work rapidly, their treatment is often reversible and they can be applied at a concentration and developmental time-point of interest, thereby circumventing problems related to lethality. In addition, gene redundancy is less of an issue as inhibitors often target related proteins, allowing simultaneous inactivation of different members of a gene family. On the other hand, the lack of specificity is often considered a drawback of pharmacological approaches, as it could come with unwanted off-target effects.

Piperonylic acid (PA) is a well-known inhibitor of CINNAMATE-4-HYDROXYLASE (C4H; (Schalk et al. 1998; Van de Wouwer et al. 2016; Desmedt et al. 2021; El Houari et al. 2021b)) and is often used to demonstrate the involvement of the phenylpropanoid pathway in distinct developmental and physiological plant processes (Naseer et al. 2012; Lee et al. 2013; Lee et al. 2019; Reyt et al. 2020). For example, we previously used PA to investigate the role of phenylpropanoid-derived lignin in phloem-mediated auxin transport (El Houari et al. 2021b). The perturbation of auxin transport in PA-treated etiolated seedlings resulted in the accumulation of adventitious roots (AR) specifically at the top part of the hypocotyl, a phenotype that could be partly complemented by restoring lignification. In this follow-up study we assess the validity of PA as an inhibitor of C4H by mapping its off-target effects.

## RESULTS

In the model plant *Arabidopsis thaliana* C4H is encoded by a single copy gene (Raes et al. 2003). As redundancy is not at play for this gene, similar effects on the phenylpropanoid pathway are to be expected for PA-treated plants and *c4h* knockout mutants. This assumption was confirmed in a previously reported experiment comparing the metabolome of etiolated mock-treated Col-0, PA-treated Col-0 and *c4h-4* mutant seedlings (El Houari et al. 2021b). The metabolite profiles of the latter two clustered closely together in a PCA plot, but separately from those of the mock-treated Col-0 samples. This indicated that genetic and pharmacological inhibition of C4H causes a similar effect on the metabolome. However, when we excluded Col-0 from the PCA analysis, the metabolic profiles obtained from *c4h-4* mutants and PA-treated seedlings resulted in the formation of two separate clusters (Fig. 1A), pinpointing at least some metabolic differences between the two conditions. The most evident explanation for this difference is the presence of PA itself, as PA was not added to the *c4h-4* mutants. A total of 398 statistically significant differentially abundant compounds were detected between the *c4h-4* mutant and PA-treated seedlings (p<0.0001). To further investigate the cause of this difference we assessed the top 15 of differential compounds between PA-treated seedlings and the *c4h-4* mutant (Table 1). All 15 compounds were present in the PA-treated samples but nearly entirely absent in the *c4h-4* mutant. Eight compounds could be characterized from this set and these were all structurally related to PA, as they were either free PA or PA-conjugates (Table 1). The highest differentially accumulating compounds were the amino acid conjugates PA-Asp and PA-Glu, with the detected quantity of PA-Asp being higher than that of all 14 other top differential compounds combined. Noteworthy was also the lower amount of free PA detected compared to its conjugates, reflecting a strong detoxification of PA by the plant.

**Table 1.** PA is conjugated by the plant. Metabolic profiling was performed for etiolated mock-treated Col-0, piperonylic acid (PA)-treated Col-0 and *c4h-4* seedlings (El Houari et al. 2021b). The table shows the detected quantities of the top accumulating compounds for PA-treated compared with *c4h-4* seedlings (n>7) for all 3 conditions (mock-treated Col-0, PA-treated Col-0 and *c4h-4* seedlings). For each of these compounds a unique number (No.), mass-to-charge ratio (*m/z*) and retention time (RT) is given.

| No. | RT | *m/z* | Name | WT | *c4h-4* | PA |
| --- | --- | --- | --- | --- | --- | --- |
| 1 | 5.64 | 280.0457 | Piperonyl aspartate | 0.00 ± 0.00 | 3.00 ± 8.21 | 9085.10 ± 2990.26 |
| 2 | 6.49 | 294.0613 | Piperonyl glutamate | 0.00 ± 0.00 | 0.00 ± 0.00 | 2176.32 ± 895.68 |
| 3 | 9.04 | 753.1494 | Unknown | 0.00 ± 0.00 | 0.00 ± 0.00 | 1001.24 ± 440.91 |
| 4 | 5.80 | 327.0711 | Piperonyl hexose | 0.00 ± 0.00 | 1.00 ± 1.54 | 710.03 ± 250.90 |
| 5 | 4.84 | 293.0772 | no MS/MS | 2.00 ± 0.59 | 0.00 ± 0.00 | 576.88 ± 238.47 |
| 6 | 5.26 | 407.0281 | Piperonyl sulfohexose | 0.00 ± 0.00 | 0.00 ± 0.00 | 531.41 ± 215.38 |
| 7 | 9.51 | 380.9547 | no MS/MS | 0.00 ± 0.00 | 0.00 ± 0.00 | 473.64 ± 161.19 |
| 8 | 5.63 | 236.0553 | Piperonyl aspartate fragment | 0.00 ± 0.00 | 0.00 ± 0.00 | 380.09 ± 128.55 |
| 9 | 5.78 | 165.019 | Piperonyl hexose | 0.00 ± 0.00 | 0.00 ± 0.00 | 349.51 ± 78.97 |
| 10 | 5.62 | 379.9698 | Unknown | 0.00 ± 0.00 | 0.00 ± 0.00 | 345.08 ± 92.89 |
| 11 | 9.52 | 165.019 | PA | 0.00 ± 0.00 | 0.00 ± 0.00 | 298.39 ± 82.70 |
| 12 | 5.77 | 363.047 | Unknown | 0.00 ± 0.00 | 0.00 ± 0.00 | 261.41 ± 95.63 |
| 13 | 4.61 | 535.1293 | PA + 2 hexoses | 0.00 ± 0.00 | 0.00 ± 0.00 | 247.16 ± 117.25 |
| 14 | 9.05 | 827.1488 | no MS/MS | 0.00 ± 0.00 | 0.00 ± 0.00 | 209.75 ± 190.79 |
| 15 | 9.50 | 615.9776 | no MS/MS | 0.00 ± 0.00 | 0.00 ± 0.00 | 204.78 ± 78.18 |

**Figure 1.**
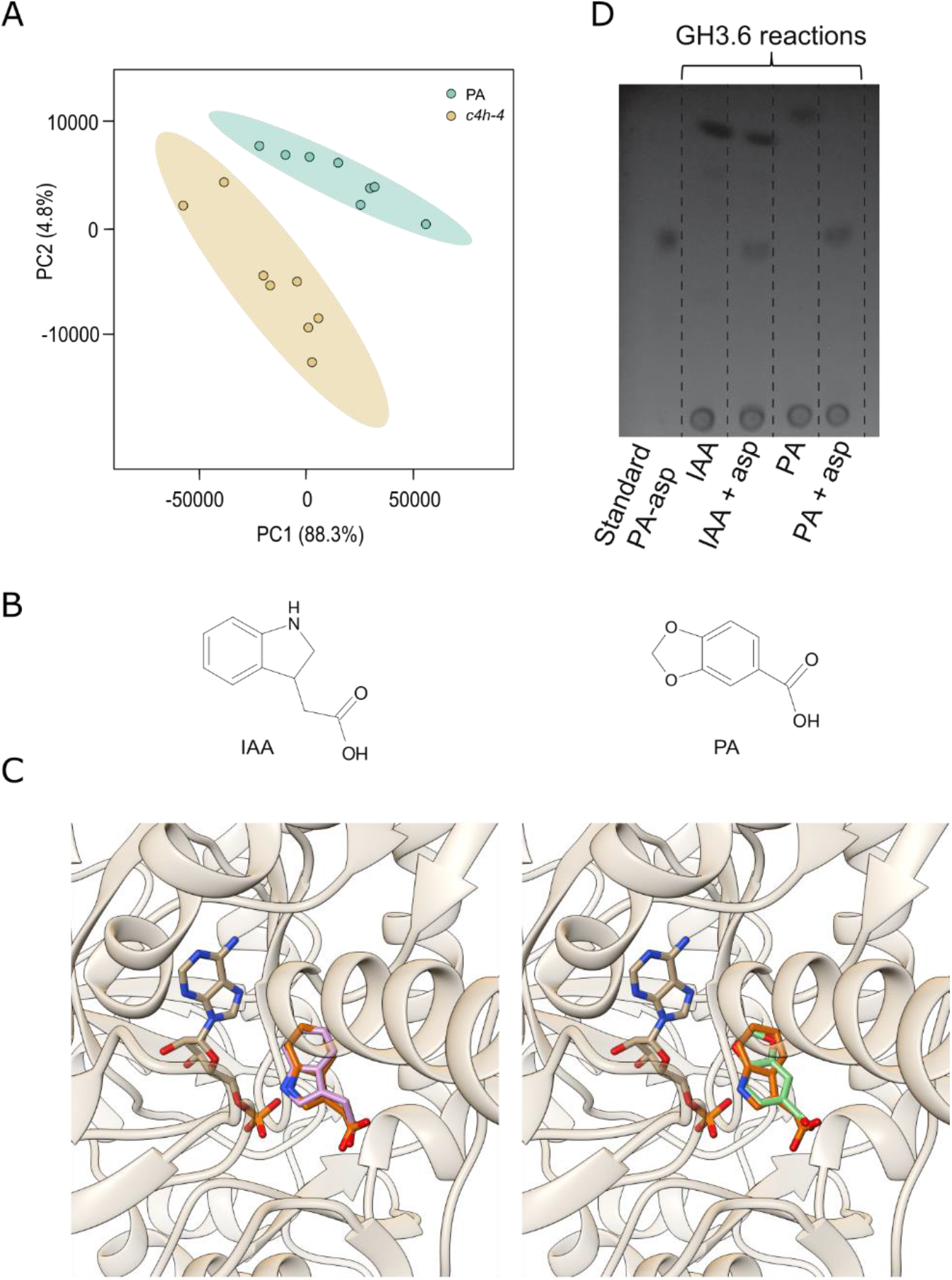
PA is recognized and conjugated by GH3.6. (A) Principal component analysis score plots for the metabolic profiles obtained by LC-MS analysis of etiolated *c4h-4* and 50 μM PA-treated Col-0 seedlings (n>7). Each data point represents eight pooled seedlings. (B) Chemical structures of indole-3-acetic acid (IAA) and piperonylic acid (PA). (C) Docking of the best possible position for IAA (left, pink) and PA (right, green) in the GH3.6 binding pocket. The experimentally determined position of IAA (orange) and adenosine monophosphate is shown for both figures. (D) TLC analysis of the products of *in vitro* enzymatic assays shows conjugation of Asp by GH3.6 to both IAA and PA.

The conjugation of metabolites to amino acids in plants is known to be conducted by the GRETCHEN-HAGEN3 (GH3) protein family (Staswick et al. 2005) and is key in the homeostasis of phytohormones and other bioactive molecules. Among these, the GH3.6-mediated conjugation of auxin (indole-3-acetic acid; IAA) to Asp, Ala, Phe and Trp is one of the best documented processes (Staswick et al. 2005). Intriguingly, PA and IAA are similar in size (166 and 175 Da, respectively) and both molecules consist of a planar aromatic carbon skeleton decorated with a carboxylic acid (Fig. 1B). Despite these similarities, both compounds have a different core carbon skeleton, PA being a benzodioxane whereas IAA is an indole. Additionally, the length of the side chains differs for both compounds, as IAA is decorated with an acetic acid and PA is decorated with a carboxylic acid. However, considering the general substrate promiscuity of the GH3s (Staswick et al. 2005), it is not unlikely that PA could also be recognized by GH3.6 as a substrate. To predict whether binding of PA to GH3.6 is possible and to estimate the likelihood of such an event, we performed a comparative *in silico* docking experiment using PA as well as IAA as substrates (Fig. 1C). As the structure of GH3.6 has not yet been solved, we did a comparative modelling using the crystal structure of GH3.5 as a template. Since GH3.6 is expected to have the same two-step catalytic mechanism as GH3.5 (Westfall et al. 2016), we retained adenosine monophosphate (AMP) within our model. The docking results for the natural ligand IAA show an excellent correspondence between the best predicted binding pose and that adopted by the substrate within GH3.5, as revealed by the crystal structure (Fig. 1C, left panel). This suggests that the binding of IAA onto GH3.6 is indeed very likely to happen via the same interactions as in GH3.5. A comparison of this result with the docked poses of PA revealed the occurrence of a bound pose identical to that of IAA (Fig. 1C, right panel) within the top 5 predicted poses for PA. This indicates that PA is a strong ligand for GH3.6. To gain empirical evidence that PA can indeed be conjugated by GH3.6, we evaluated the conjugation of PA by GH3.6 *in vitro* (Fig. 1D). As a positive control we provided GH3.6 with both IAA and Asp, which resulted in the formation of IAA-Asp. Supplying GH3.6 with both PA and Asp resulted in the formation of the PA-Asp conjugation product, demonstrating that PA can indeed be conjugated to Asp by GH3.6 *in vitro*.

Having shown that GH3.6 conjugates PA to Asp, we speculated that a major increase in PA levels could overload the catabolic machinery of the plant and thus obstruct the conjugation of IAA. To verify this model, we assessed whether PA-treatment could indeed inhibit or slow down IAA conjugation. For this purpose, we implemented a cellular auxin conjugation assay, in which BY2 cell cultures are fed with the radiolabeled synthetic auxin analog [^3^H]NAA. When supplemented, NAA enters the cell passively but is exported actively out of the cell (Delbarre et al. 1996). Inside the cell, the radiolabeled NAA is conjugated by a range of catabolic enzymes, including GH3.6. This conjugation makes NAA unavailable for auxin exporters and traps the signal inside the cell. We hypothesized that should PA interfere with IAA conjugation, the NAA entering the cell would not be conjugated and thus remain available for export, resulting in a lower end-point signal compared with mock-treated samples. As expected, treatment of the cell cultures with only [^3^H]NAA resulted in a steady increase in signal over time, as a fraction of [^3^H]NAA is conjugated and can therefore not be exported (Fig. 2A). Upon co-treatment of the cell-cultures with [^3^H]NAA and PA, the intracellular level of [^3^H]NAA quickly reached a plateau, with final [^3^H]NAA levels significantly lower compared to those of mock-treated samples (Fig. 2A). These results further indicate that PA-treatment indeed impedes auxin conjugation.

**Figure 2.**
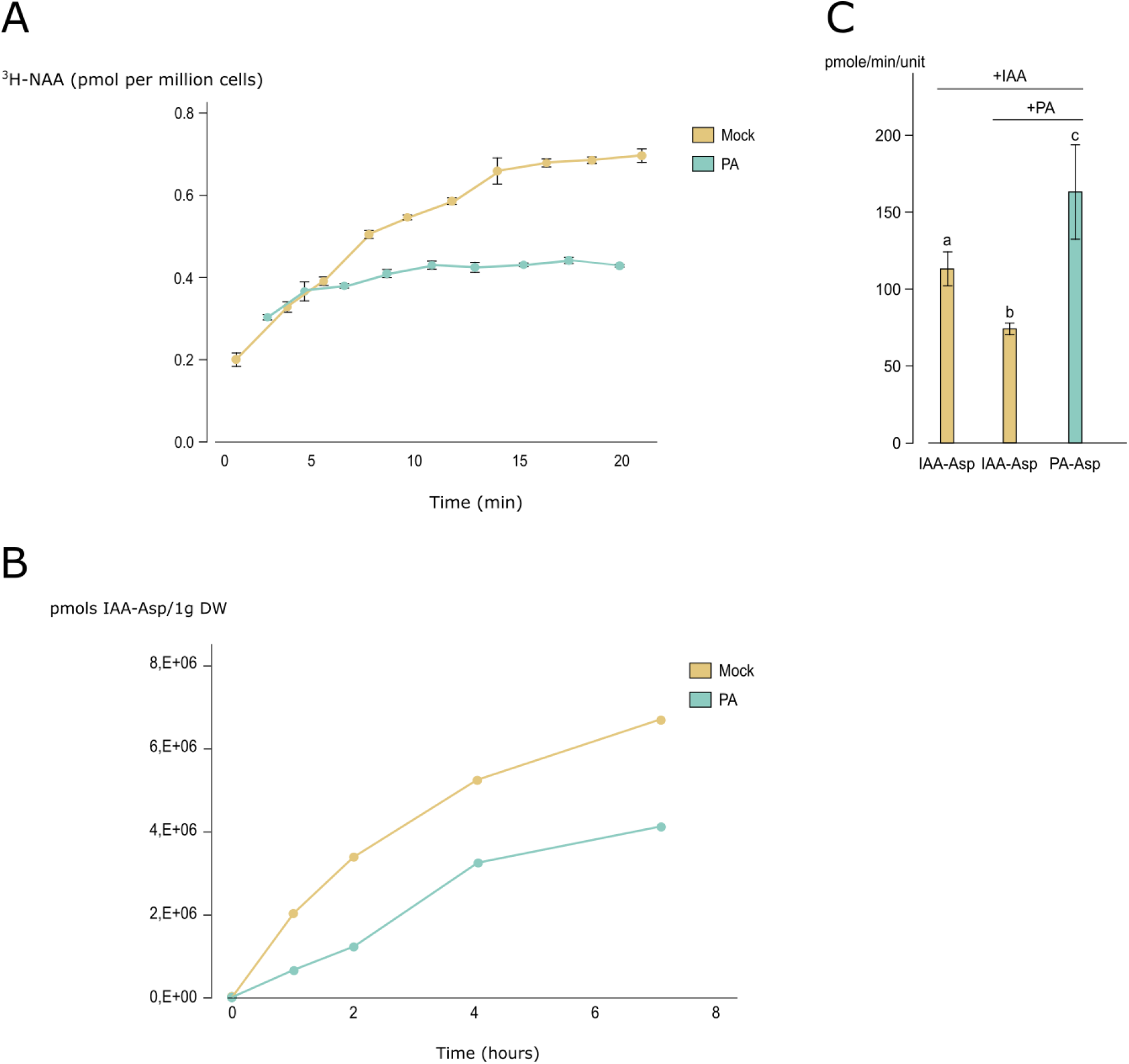
PA treatment slows down the conjugation of IAA to Asp by GH3.6. (A) Cellular auxin conjugation assay in BY-2 cells using radiolabeled [^3^H]NAA over time upon treatment with or without PA (n=4). Error bars represent standard error. (B) Quantification of IAA-Asp in BY-2 cells over time upon treatment with IAA and with or without PA. (C) Quantification of the products IAA-Asp (yellow) and PA-Asp (blue) upon supplying GH3.6 *in vitro* with IAA and/or PA. Error bars represent 95% confidence intervals. Letters a-c are given to distinguish statistically significant values (P<0.01; (a– c) ANOVA; Tukey’s honestly significant difference test).

These results are however not conclusive for a PA-mediated obstruction of the conjugation of IAA to Asp in the cell, as NAA is a synthetic analog of IAA and as we did not specifically assess conjugation to Asp. Therefore, we next verified whether PA-treatment could interfere with the conjugation of IAA to Asp in a cellular context (Fig. 2B). For this, IAA-Asp concentrations were followed over time in BY2 cell cultures upon addition of 10 µM IAA with or without PA. As anticipated, the concentration of IAA-Asp produced over time was significantly lower upon addition of PA, confirming the hypothesis that PA slows down the conjugation of IAA to Asp (Fig. 2B), likely by competing for the same enzymes. To obtain conclusive evidence that PA interferes with the conjugation of IAA to Asp by GH3.6, we quantified the levels of IAA-Asp formed upon supplying GH3.6 *in vitro* with either IAA and Asp or IAA, Asp and PA (Fig. 2C). These data showed a significant reduction in the levels of IAA-Asp formed upon co-treatment with PA. In addition, the levels of PA-Asp formed were significantly higher than those of IAA-Asp. These results show that PA effectively slows down the catabolism of IAA to IAA-Asp by GH3.6.

So far, we examined the involvement of GH3.6 in the conjugation of PA. To assess an involvement of the other GH3s in PA-conjugation, we quantified the shift in expression of IAA-conjugating *GH3* genes in mock- or PA-treated seedlings (Fig. 3A; (Staswick et al. 2005)). Of the six *GH3* genes tested, five showed a significant upregulation upon PA-treatment (i.e. *GH3*.*1, GH3*.*2, GH3*.*3, GH3*.*5* and *GH3*.*6*), with only *GH3*.*4* expression not significantly changed. These results point towards a strong *GH3*-mediated response in PA-treated plants. Treatment with PA was previously shown to strongly induce AR growth in seedlings and to do this specifically at the top part of the hypocotyl (El Houari et al. 2021b). We therefore assessed whether the interference of PA with the conjugation of IAA by GH3s would contribute to this phenotype. To do so, we compared the AR growth of a sextuple *gh3* mutant defunct for the same *GH3* genes whose expression we previously assessed (*gh3*.*1,2,3,4,5,6*; Fig. 3B) to AR growth in mock- and PA-treated Col-0 seedlings. ARs were quantified while also considering their localization on the hypocotyl, being either at the top third part or the bottom two thirds part. As previously described, PA-treated Col-0 seedlings displayed a strong increase in total ARs compared to the mock-treated Col-0 plants and this increase was specifically situated at the top part of the hypocotyl (El Houari et al. 2021b). Correspondingly, the *gh3* sextuple mutants also showed a strong induction of AR compared to the mock-treated Col-0 plants (Fig. 3B), albeit along the entirety of the hypocotyl. These results thus demonstrate that prohibiting GH3-mediated conjugation of IAA upon PA-treatment could indeed contribute to an overall increase in AR growth proliferation.

**Figure 3.**
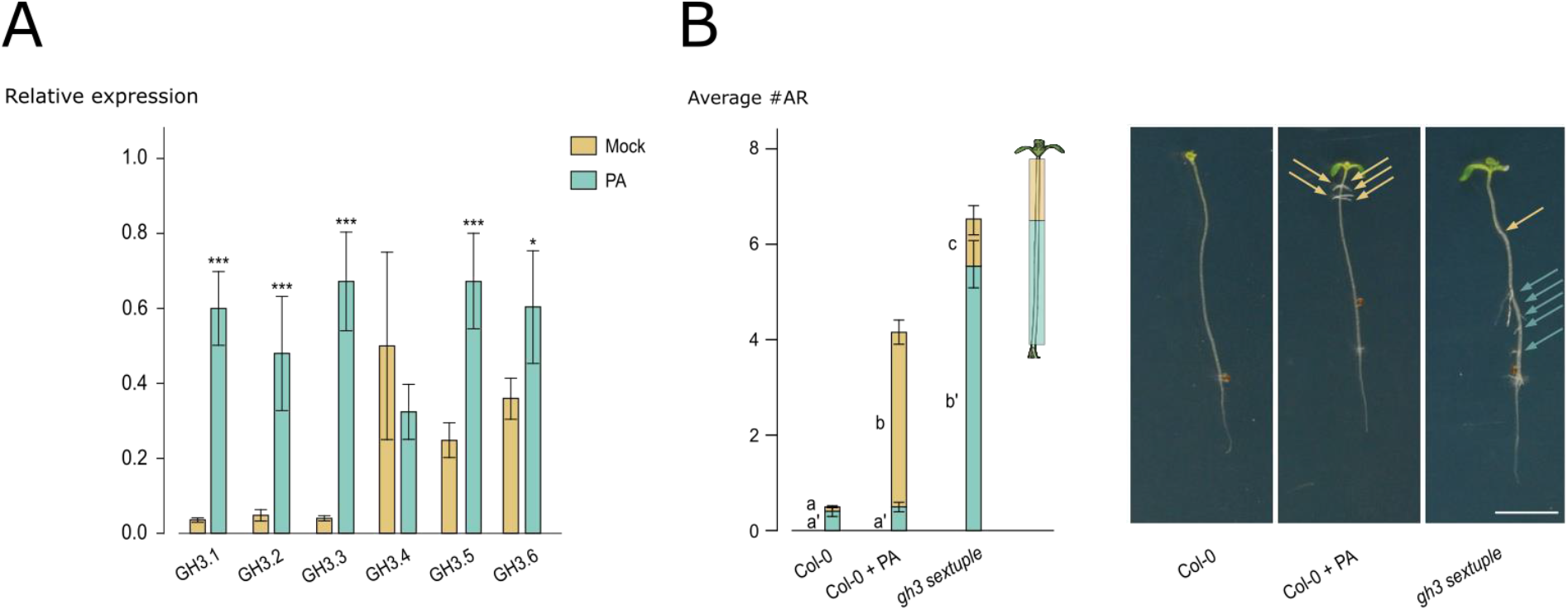
Obstruction of GH3-mediated auxin catabolism results in increased adventitious rooting. (A) Expression levels of *GH3*.*1-6* in mock-treated and PA-treated etiolated seedlings (n=9). Error bars represent 95% confidence intervals. Asterisks indicate significant differences compared to the corresponding mock-treatment (*, P<0.01; **, P<0.001; ***, P<0.0001; Student’s t-test) (B) Average number of adventitious roots (ARs) of etiolated mock-treated Col-0, 50 μM PA-treated Col-0 and *gh3* sextuple mutant seedlings (n>20). Yellow coloration, top third part of the hypocotyl; blue coloration, lower two-thirds part of the hypocotyl. On the right, a representative seedling is presented for each of the conditions. Bar=1cm. Yellow arrow, ARs located at the top third part of the hypocotyl; blue arrow, ARs located at the bottom two-thirds part of the hypocotyl. Error bars represent 95% confidence intervals. Letters a-c are given to distinguish statistically significant values (P<0.01; GEE model).

## DISCUSSION

Plants make extensive use of small compounds to steer their growth and development. As these bioactive compounds can easily negatively affect plant growth when mislocalized or when over or under abundant, their availability is under tight control. Accordingly, plants are equipped with a range of enzymes that mediate the conjugation and/or sequestration of these compounds, such as UDP-glycosyltransferases (UGTs), glutathione-S-transferases (GSTs) and amido synthetases (Schröder and Collins 2002; Casanova-Sáez et al. 2021). For example, the glycosylation of several phenylpropanoids allows for the regulation of their endogenous levels via sequestration to the vacuole (Dima et al. 2015; Le Roy et al. 2016), a mechanism which is proposed to mitigate the toxicity of bioactive phenylpropanoid accumulation (Le Roy et al. 2016; Vanholme et al. 2019; El Houari et al. 2021a; Steenackers et al. 2019). Such conjugating enzymes tend to have large substrate promiscuities and can act both on endogenous compounds as well as compounds that are exogenous to the plant (Staswick et al. 2005; Mateo-Bonmatei and Ljung 2021; Aoi et al. 2020). Consequently, when exogenous compounds are supplied in excess, their inactivation could overwhelm the pool of catabolic enzymes and jeopardize the homeostasis of endogenous bioactive compounds.

Here, we demonstrate that piperonylic acid (PA), an inhibitor of CINNAMATE-4-HYDROXYLASE (C4H), is recognized by GRETCHEN HAGEN 3.6 (GH3.6), an enzyme known to be involved in the conjugation of amino acids to several molecules. One of the best-studied substrates of GH3.6 is the phytohormone indole-3-acetic acid (IAA). We show that excessive PA treatment effectively slows down auxin conjugation, resulting in an increase in the intracellular levels of free auxin. Specifically, we show that PA can slow down conjugation of auxin to Asp by GH3.6, hereby likely contributing to visible phenotypes. Although we focused on GH3.6, it is likely that PA can also be recognized by other GH3 enzymes and can interfere with their normal cellular activity. In addition, the perturbation should not be limited to the amino acid conjugating enzymes. Glucosyl conjugation products of PA were also highly accumulating in PA-treated seedlings (Table 1), indicating that also the conjugation of auxin to sugars by UDP-glycosyltransferases (UGTs) can be impaired.

PA-treated plants show an accumulation of AR and these AR are typically located in the top part of the hypocotyl (Fig. 3B). This was shown to be caused by a perturbation in auxin transport upon inhibition of C4H by PA (El Houari et al. 2021b). Correspondingly, the *gh3* sextuple mutant also showed a strong induction of AR compared to the mock-treated Col-0 plants. However, in contrast to PA-treated seedlings, the AR growth in the *gh3* sextuple mutant was increased along the entire hypocotyl instead of specifically at the top third part. Importantly, *c4h-4* mutant seedlings also showed an increase of AR specifically in the top part of the hypocotyl, despite not being treated with PA (El Houari et al. 2021b). Together, these results seem to indicate that prohibiting GH3-mediated conjugation of IAA upon PA-treatment could indeed contribute to an overall increase in AR growth proliferation. However, the specific apical induction of AR is not caused by the interference of PA with IAA conjugation *per se*. Rather, it is likely to be caused by the inhibition of C4H and the consequential perturbation of auxin transport, as previously described (El Houari et al. 2021b). The increase in AR observed upon knocking out *GH3s* could also explain for some slight phenotypic differences between PA-treated plants and the *c4h-4* mutant. In the *c4h-4* mutant, the auxin redistribution in the hypocotyl upon inhibition of auxin transport goes along with a decrease in AR at the bottom part of the hypocotyl (El Houari et al. 2021b). In contrast, PA-treated seedlings rarely show such decrease in the number of ARs in this region, regardless of the PA concentrations used. This phenotypic difference can be explained by the obstruction of auxin catabolism upon treatment with PA. The resulting higher levels in free auxin counteract the depletion in auxin at the bottom part of the hypocotyl caused by a perturbed auxin transport. This results in a higher number of AR in this region upon PA-treatment compared with the *c4h-4* mutant. This hypothesis is consistent with the large increase in ARs observed at the bottom part of the hypocotyl in the *gh3* sextuple mutant (Fig. 3B).

The conjugation of endogenous plant hormones by GH3s and UGTs has not only been described for auxin but also for other phytohormones, such as jasmonate and salicylic acid (Zhang et al. 2007; Ding et al. 2008; Westfall et al. 2016; Casanova-Sáez and Voß 2019). Therefore, PA-treatment could influence the endogenous levels of not only auxin but several other bioactive molecules, thereby indirectly affecting a large array of biological processes. Also, and importantly, treatment with other exogenous compounds will likely also obstruct the metabolism of endogenous molecules in the same manner. Therefore, other chemical inhibitors could, analogously to PA, influence phytohormonal homeostasis by hijacking the plant conjugation machinery. As a consequence, the transcriptome, proteome and metabolome might be altered by such treatment in an indirect manner, causing erroneous conclusions to be drawn. We therefore advise to take into account and assess the catabolism of the exogenous compound by the plant, as this could give valuable insight into possible off-target effects caused by the implemented compound and prohibit confusing primary with secondary effects.

## METHODS

### Plant material, transgenic lines, chemicals and growth conditions

*Arabidopsis thaliana* of the Col-0 ecotype was used for all analyses. The *c4h-4* mutant (GK-753B06; (Kleinboelting et al. 2012)) was obtained from the NASC institute. Seeds were vapor-phase sterilized and plants grown on ½ Murashige & Skoog (MS) medium (pH 5.7) containing 2.15 g MS basal salt mixture powder (Duchefa), 10 g sucrose, 0.5 g MES monohydrate, 8 g plant tissue culture agar per liter. When relevant, the medium was supplemented with either dimethyl sulfoxide (DMSO) as a mock treatment or piperonylic acid (PA; Sigma Aldrich). This compound was prepared as a stock solution in DMSO and was added to the autoclaved medium before pouring the plates. Seeds were stratified via a 2-d cold treatment. Adventitious rooting induction was performed as described previously (El Houari et al. 2021b).

### Phenotyping

Adventitious rooting was analyzed as described in El Houari et al., 2021. Subsequent statistical analyses of rooting phenotypes were also performed as described in El Houari et al., 2021.

### Metabolic profiling and analysis

The data used for the metabolic profiling was obtained from El Houari et al., 2021. To detect significant differential metabolites between the *c4h-4* and PA-treated seedlings we applied several criteria: (1) Peaks should be present in all samples of at least one out of two conditions; (2) Student’s t-test *P* < 0.0001; (3) average normalized abundance should be higher than 100 counts in at least one out of two conditions; (4) there should be at least a 100-fold difference in peak area between the two conditions. From this set, the 15 most abundant peaks were selected and sorted by detected quantities in PA-treated samples. Annotation of compounds matching these criteria was based on accurate *m/z*, isotope distribution, and tandem mass spectrometry (MS/MS) similarities. Compounds were structurally elucidated based on similarity of their MS/MS spectra with commercially available standards and previously identified metabolites that were already described in the literature.

### Homology modelling and docking

To create a putative structure of GH3.6 Modeller 10.1 was used (Šali and Blundell 1993). Chain B of the crystal structure of AtGH3.5 was selected as template, since it has a sequence identity of 91% with AtGH3.6 on an alignment over 573 residues out of 612. Note that of the 39 non-aligned residues, all but 14 were found at the termini of the protein, where short disordered loops were not crystallized. 64 different initial models were built, performing a slow annealing stage twice on each one. Each model was then refined 16 independent times, specifically targeting the non-aligned region between R376 and A389 to predict its folded state using loop refinement (Fiser and Do 2000). In all the resulting 1024 models, the presence of AMP within the binding site was retained. To identify the best model, each was scored according to a high-resolution version of the Discrete Optimized Protein Energy, or DOPE-HR (Shen and Sali 2006), and the model with the best score that did not exhibit structural clashes was chosen. All docking runs were performed with Autodock Vina (Vina 2010). A search space of 7400 cubic Å (20×20×18.5) centered on the binding site (x, y and z coordinates -2.04, 101.2 and 94.73, respectively) was set and a search exhaustiveness of 128 was used. Ligand files were drawn and energy-minimized in Avogadro2 (Hanwell et al. 2012). Ligand files and model were prepared for docking using AutoDockTools (Morris et al. 2009). Docked poses were evaluated visually using IAA as the reference. All visualizations were produced using UCSF Chimera (Pettersen et al. 2004).

### Enzyme assays

IAA conjugation assays were done using GH3.6-GST fusion protein produced in *E. coli* as previously described (Staswick et al., 2005). For kinetic reactions enzyme was released from GST beads using reduced glutathione. Qualitative analysis reactions (Fig. 1D) were performed for 16 h at 23°C in 50 mM Tris-HCl, pH 8.6, 1 mM MgCl2, 1 mM ATP, 1 mM DTT, and 2 mM Asp. Either IAA (1 mM) or PA (10 mM) was included in each reaction. Reactions were analyzed on silica gel 60 F260 plates developed in chloroform:ethyl acetate:formic acid (35:55:10, v/v) and then stained with vanillin reagent (6% vanillin [w/v], 1% sulfuric acid [v/v] in ethanol). Kinetic experiment assays (Fig. 2C) were carried out as described for JA-Ile conjugation (Suza and Staswick 2008), substituting PA, IAA and Asp (1 mM each) as the substrates with the GH3.6 enzyme. Results were extrapolated over a linear range that included assay timepoints of 2,5,8 and 10 min. Reaction products were quantified by GC/MS using ^13^C_6_ IAA-Asp as an internal standard for IAA and using a linear standard curve for PA-Asp, the latter synthesized and purified as previously described for JA conjugates (Staswick and Tiryaki 2004).

### Cellular auxin conjugation assays

Assays were performed according to (Petrášek et al. 2003). Auxin accumulation was measured in tobacco BY-2 cells (*Nicotiana tabacum* L. cv. Bright Yellow 2; (Nagata et al. 1992)) 48 hours after subcultivation. Cultivation medium was removed by filtration on 20 μm mesh nylon filters and cells were resuspended in uptake buffer (20 mM MES, 10 mM sucrose, 0.5 mM CaSO4, pH adjusted to 5.7 with KOH) and equilibrated for 45 minutes on the orbital shaker at 27 °C in darkness. Cells were then collected by filtration, resuspended in fresh uptake buffer and incubated for another 90 minutes under the same conditions. Radiolabelled auxin ([^3^H]naphthalene-1-acetic acid (^3^H-NAA); specific radioactivity 20 Ci/mmol; American Radiolabeled Chemicals, ARC Inc., St. Louis, MO, USA) was added to the cell suspension to the final concentration of 2 nM. 0.5 ml aliquots of cell suspension (density 7×10^5^ cells×ml^-1^) were sampled and accumulation of auxin was terminated by rapid filtration under reduced pressure on cellulose filters. Samples with filters were transferred into scintillation vials, extracted with ethanol for 30 minutes and radioactivity was determined by liquid scintillation counting (Packard Tri-Carb 4910TR scintillation counter, Packard Instrument Co., Meridien, CT, USA). Counting efficiency was determined by automatic external standardization and counts were corrected for quenching automatically. Counts were corrected for remaining surface radioactivity by subtracting counts of aliquots collected immediately after addition of ^3^H-NAA. Piperonylic acid and solvent control (DMSO) were applied 1 minute after the start of the experiment. Recorded accumulation values were recalculated to pmol/1 million cells.

### Cellular IAA-Asp conjugation assays

Cellular auxin metabolites were determined according to (Dobrev et al. 2017) in tobacco BY-2 cells (*Nicotiana tabacum* L. cv. Bright Yellow 2; (Nagata et al. 1992)) supplied with 10 μM IAA and 50 μM piperonylic acid 48 hours after subcultivation. The homogenized samples (about 0.1 g fresh weight (FW) were incubated in 0.5 ml extraction buffer (methanol : formic acid : water, 15 : 1 : 4, v/v) with internal standards (10 pmol) for 1 h at -20°C. After centrifugation (20 000 g, 20min, 4°C), the supernatant was collected, and the pellet was re-extracted with additional extraction buffer (30min, -20°C). Pooled supernatants were evaporated in a vacuum concentrator (Alpha RVC; Christ, Osterode am Harz, Germany), re-dissolved in 0.5ml of 1M formic acid and applied to an SPE column (Oasis-MCX; Waters, Milford, MA, USA). The auxin fraction was eluted with 0.35M NH4OH in 70% methanol, evaporated to dryness and dissolved in 30 μl 5% methanol. An aliquot (3 μl) was analyzed on an Ultimate 3000 HPLC device (Dionex, Sunnyvale, CA, USA) coupled to MS (3200 Q TRAP; Applied Biosystems, Foster City, CA, USA) as described previously (Dobrev and Vankova 2012; Dobrev et al. 2017).

### RNA isolation and qRT-PCR analysis

Total RNA was isolated from etiolated seedlings grown according to El Houari et al., 2021 with TriZol (Invitrogen), purified with the RNeasy Plant Mini Kit (Qiagen) and treated with DNase I (Promega). Complementary DNA (cDNA) was prepared with the iScript cDNA Synthesis Kit (Bio-Rad) according to the manufacturer’s instructions. Relative transcript abundancies were determined using the Roche LightCycler 480 and the LC480 SYBR Green I Master Kit (Roche Diagnostics). The resulting cycle threshold values were converted into relative expression values using the second derivative maximum method and *ACTIN2, ACTIN7* and *UBIQUITIN10* were used as reference genes for normalization. All experiments were performed in three biological replicates (∼10 seedlings per replicate), each with three technical replicates. The primer sequences are listed in Supplemental Table S1.

## ACKNOWLEDGEMENTS

This work was supported by the Fonds voor Wetenschappelijk Onderzoek – Vlaanderen (FWO) through project numbers G008116N and 3G038719 and also by personal grants to IE. by FWO (1S04020N; V414521N) and EMBO (STF-8658). PS was supported by the University of Nebraska by Agricultural Research Division, funded in part by the USDA. CIDG acknowledges support from UKRI under Future Leaders Fellowship grant number MD/T020652/1. P.K. acknowledges support from the Ministry of Education, Youth, and Sports of the Czech Republic (project no. CZ.02.1.01/0.0/0.0/16_019/0000738, EU Operational Programme “Research, development and education and Centre for Plant Experimental Biology”). We would like to thank the VIB Metabolomics Core Facility and Geert Goeminne for processing of the LC-MS samples. We would like to thank Ruben Vanholme and Kris Morreel for help with the analysis of the metabolomics data. We would also like to thank Karel Spruyt for help with the photo.

## FIGURE AND TABLES

**Supplemental table 1.**
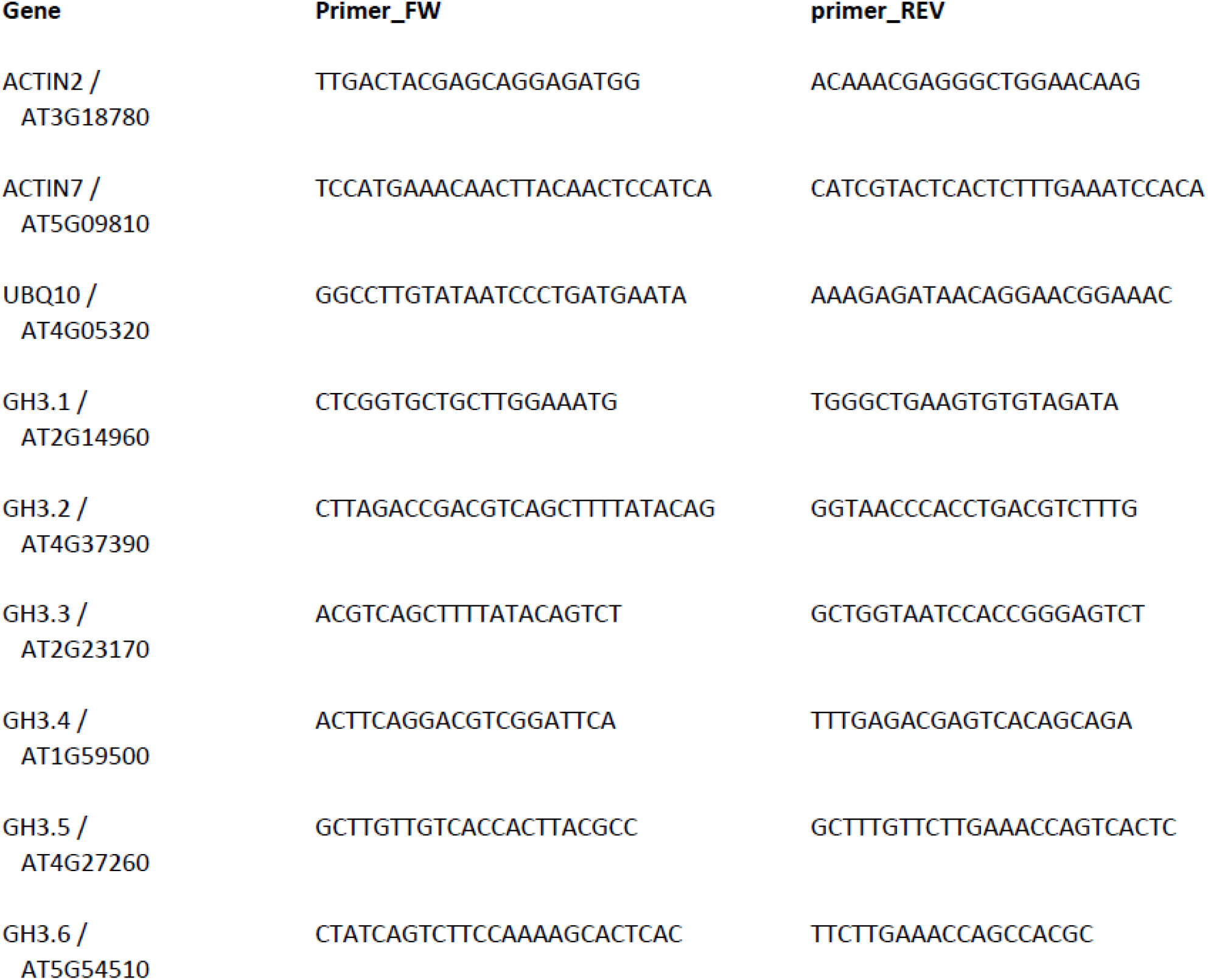
List of primers used for qRT-PCR analysis.

